# A novel glutathione peroxidase-based biosensor disentangles differential subcellular accumulation of H2O2 and lipid hydroperoxides

**DOI:** 10.1101/2024.01.18.576236

**Authors:** Marino Exposito-Rodriguez, Brandon Reeder, Greg N. Brooke, Michael A. Hough, Philippe P. Laissue, Philip M. Mullineaux

**Affiliations:** School of Life Sciences, University of Essex, Wivenhoe Park, Colchester CO4 3SQ, UK; Diamond Light Source Ltd, Harwell Science and Innovation Campus, Didcot OX11 0DE, UK

**Keywords:** Biosensors, plants, live imaging, hydrogen peroxide, lipid hydroperoxides, (phospholipid) fatty acid hydroperoxides, retrograde signaling, oxidative stress, *Nicotiana benthamiana*, glutathione peroxidase

## Abstract

Hydrogen peroxide (H_2_O_2_) and lipid hydroperoxides (LOOH) are initiators and transducers of inter- and intra-cellular signaling in response to diverse environmental, pathological and developmental cues. The accumulation of both H_2_O_2_ and LOOH is often temporally and spatially coincident in tissues, but it is unknown if this coincidence extends to subcellular compartments. If distinct accumulation of different peroxides occurs at this smaller spatial scale, then it would be an important factor in signaling specificity. Fusion of the redox-sensitive (ro)GFP2 to the *Saccharomyces cerevisiae* (yeast) OXIDANT RECEPTOR PEROXIDASE1 (ORP1), also known as GLUTATHIONE PEROXIDASE3 (GPX3), created a now widely used biosensor that is assumed to detect H_2_O_2_ *in vivo.* This is despite monomeric GPX enzymes, such as ORP1/GPX3, possessing wide peroxide substrate specificities. Consequently, we confirmed *in vitro* that roGFP2-ORP1 is not only oxidized by H_2_O_2_, but also by phospholipid fatty acid peroxides generated in lecithin-derived liposomes by lipoxygenase-catalyzed peroxidation. This led us to doubt that roGFP2-ORP1 *in vivo* is specific for H_2_O_2_. To address this issue of peroxide specificity, we constructed a modified biosensor called roGFP2-synORP1. This version has greatly diminished reactivity towards phospholipid fatty acid peroxides but retains high sensitivity for H_2_O_2_. These two roGFP2-based biosensors, targeted to chloroplasts, cytosol and the nucleus, were quantitatively imaged in parallel in *Nicotiana benthamiana* abaxial epidermal cells experiencing high light- and herbicide-induced photo-oxidative stress. From differential patterns of oxidation of these probes, we inferred that the chloroplasts accumulated both peroxide types. In contrast, LOOH and H_2_O_2_ accumulated exclusively in the cytosol and nucleus respectively. Therefore, this suggests that the signalling networks initiated by different peroxides will have a distinct spatial component.

## INTRODUCTION

All aerobic organisms generate reactive oxygen species (ROS) as an unavoidable by-product of the metabolism of ground state molecular oxygen (O_2_) that occurs during respiration, photosynthesis and various metabolic reactions (Halliwell & Gutteridge, 2015; Jones and Sies 2015; Hasanuzzaman et al., 2020; Dogra and Kim 2020). ROS is a collective term for oxygen radicals (e.g. the superoxide anion) and non-radical species (e.g., hydrogen peroxide (H_2_O_2_) and singlet oxygen (^1^O_2_)). ROS are reduced or quenched by a redundant system of antioxidants and antioxidant enzymes (Buettner et al 2013; Halliwell & Gutteridge, 2015; Hasanuzzaman et al 2020). The excessive accumulation of ROS in cells causes oxidative stress (Halliwell & Gutterridge, 2015; Mullineaux and Baker 2010). At lower or more localized levels, ROS engage in intra- and extracellular signaling pathways that elicit adjustments to metabolism and growth and consequently determines the degree of susceptibility to environmental stress and disease (Buettner et al 2013; Jones and Sies 2015; Sies 2017; Mullineaux and Baker 2010; Evans et al 2016; Smirnoff and Arnaud 2019; Dogra and Kim 2020; Mittler et al 2022).

H_2_O_2_ is a potent initiator and transducer of intracellular signals (D’Autreaux and Toledano, 2007; Jones and Sies 2015; Sies 2017; Smirnoff & Arnaud 2019; Mittler et al 2022). The accumulation of H_2_O_2_ in subcellular microdomains may be an important feature in the specificity of its signaling functions (Ushio-Fukai 2009; Sies 2017; Exposito-Rodriguez et al 2017; Smirnoff and Arnaud 2019; Wu et al 2020; Niemeyer et al 2021; Bi et al 2022). In plants, H_2_O_2_ levels are associated with the induction of resistance to infection by biotrophic pathogens, acclimation to fluctuating environmental conditions, the growth of secondary root hairs and pollen tubes, and the lignification of cell walls (Chaouch et al., 2010; Wu et al 2010; Kadota et al 2014; Zhang et al 2015; Laitinen et al 2017; Rasool et al 2017; Evans et al 2016; Smirnoff and Arnaud, 2019; Wu et al 2020; Bi et al 2022; Zhang et al 2023).

^1^O_2_ is generated from the reaction of O_2_ with photosensitized pigments, such as triplet state chlorophyll P680 (^3^Chl) and phenylphenalenone phytoalexins (Thoma et al 2003; Lazzaro et al 2004; Rutherford and Krieger-Liszkay 2001; Flors et al 2006; Dogra et al 2018). Under diverse stress conditions, cells also accumulate a range of organic hydroperoxides formed non-enzymically by the reaction of ^1^O_2_ or free radicals such as the hydroxyl radical, with fatty acids, sterols and terpenes (Halliwell & Gutteridge, 2015; Sousa et al 2017; Tsubone et al 2019; Adjemian et al 2020; Muñoz and Munné-Bosch 2020). Organic hydroperoxides also cause the accumulation of reactive carbonyl and aldehyde species, which interact with downstream programmed cell death (PCD) and hormone signaling networks (Thoma et al 2003; Biswas and Mano 2015; Biswas et al 2019; Adjemian et al 2020; Muñoz and Munné-Bosch 2020; Yuan et al 2021). The formation of ^1^O_2_ has clear signaling consequences, especially PCD and induction of resistance to photo-oxidative stress (Kim et al., 2012; Ramel et al., 2013; Dogra et al 2018; Munoz and Munné-Bosch 2020). In particular, the (poly)unsaturated fatty acid peroxides (Triantaphyllides et al 2008; Tsubone et al 2019) are biosynthetic precursors for the stress defense hormone jasmonic acid (Wasternack and Feussner 2018, Ramel et al 2013).

To further understand these fundamentally pivotal roles of ROS it is important that the subcellular distribution and dynamics of H_2_O_2_ and organic peroxides be determined. In this respect, the development of genetically encoded fluorescent protein (FP) biosensors for the detection of H_2_O_2_ has proved to be seminal (Belousov et al 2006; Bilan et al 2012; Jones and Sies 2015; Morgan et al 2016; Carmona et al 2019). In plants and algae, progress on understanding spatial and temporal dynamics of H_2_O_2_ has been made using the FP biosensors HyPer and redox (ro)GFP2-TsaΔC_R_ (Costa et al 2010; Hernandez-Barrera et al 2015; Caplan et al 2015; Exposito-Rodriguez et al 2017; Calero-Muñoz et al 2019; Niemeyer et al 2021; Ugalde et al 2021b). In this study, we have used the redox-relay proximity probe based on roGFP2 connected by a 32-amino acid peptide to the *Saccharomyces cerevisiae* (yeast) OXIDANT RECEPTOR PEROXIDASE1 (ORP1; Fig. 1A; Delaunay et al 2002; Gutscher et al 2009; Schwarzländer et al 2016; Nietzel et al 2019; Ugalde et al 2021a). Changes in the redox state of this biosensor can be followed by dual excitation of the FP moiety at 405 nm (F405) and 480 nm (F488) and measuring their emission at 530 nm. The F405/F488 emission ratio, proportional to the redox state of the probe, is independent of its concentration (Belousov et al 2006; Gutscher et al 2009; Morgan et al 2016; Schwarzlander et al 2016). This is an essential property of roGFP2-ORP1 for its use *in vivo*, allowing relative levels of peroxide to be inferred, although determination of absolute levels *in vivo* is currently not feasible (Schwarzlander et al 2016).

**Figure 1.**
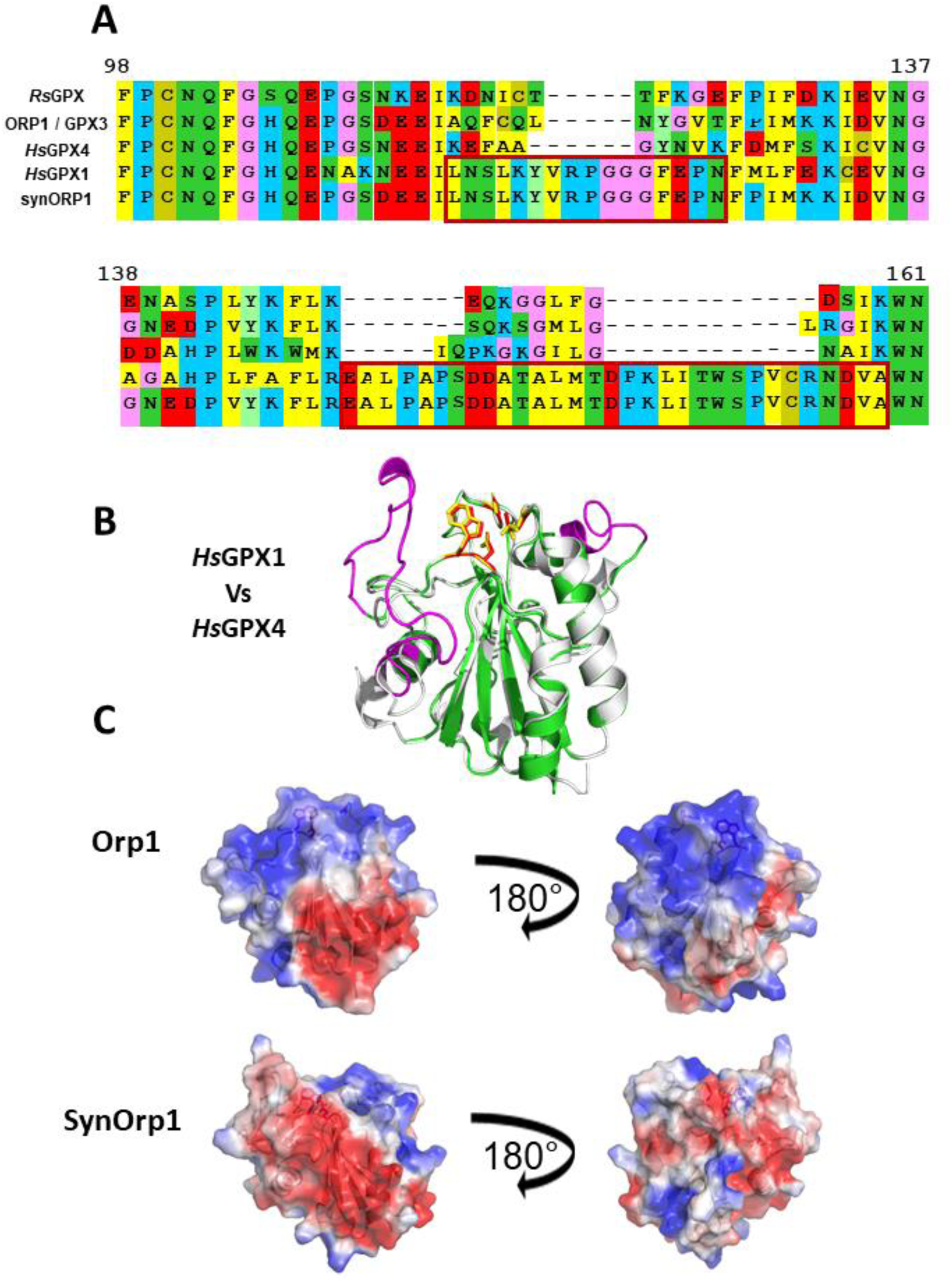
Alignment and comparison of the overall protein structures of GPX enzymes with differing peroxide substrate specificities. (A) Multiple alignment of GPXs of *Raphanus sativus* (RsGPX; XP_018476980)*, Saccharomyces cerevisiae* (ORP1; NP_012303)*, Homo sapiens* (*Hs*GPX1; AAH11836 and *Hs*GPX4; AAH11836) and a synthetic ORP1 (synORP1; this paper). After initial structural comparisons, the sequences from *Hs*GPX1 (in the rectangles) were designed into a synthetic ORP1 coding sequence, to create synORP1. Only residues 98-161 (counted from RsGPX) of each GPX are shown. (B) Structural alignment of *Hs*GPX1 (PDB 2F8A) and *Hs*GPX4 (PDB 2OBI). Both structures are depicted in cartoon representation. The *Hs*GPX1 is colored white, and the *Hs*GPX4 is colored green. The sequences conserved in *Hs*GPX1 but absent from aligned *Hs*GPX4 are colored in magenta. The catalytic centers are marked in red and yellow. (C) Comparison of the surface electrostatic potentials was calculated using the X-ray crystal structure of ORP1 (PDB 3CMI) and simulated by homology modelling of synORP1 using ProMod3 (see Methods). Two representations of each structure are shown rotated by 180 degrees. The electrostatic gradient is shown from the most negatively charged surface (red color;-2 KT·e^−1^) to the most positively charged surface (blue color;2 KT·e^−1^). Surface electrostatic potentials of *Rs*GPX, *Hs*GPX1 and *Hs*GPX4 can be found in Supplemental Figure 1.

In yeast, ORP1 is central to the operation of a H_2_O_2_-activated yAP1 transcription factor regulon (Delaunay et al 2002; D’Autreaux and Toledano, 2007; Belswinder et al 2015). While the peroxidase activity of ORP1 is not essential for its H_2_O_2_ signal transduction role in yeast (Delaunay et al 2002; Avery et al 2004), it is indispensable when ORP1 is part of a redox relay peroxide biosensor (Gutscher et al 2009; Schwarzlander et al 2016). ORP1 is also called GLUTATHIONE PEROXIDASE3 (GPX3; Avery et al 2004), catalyzing not only the thioredoxin-/glutathione- (GSH) dependent reduction of H_2_O_2_ but also tertiary butyl hydroperoxide (TBP) and the phospholipid phosphatidylcholine hydroperoxide (Avery et al 2004; Gutscher et al 2009). Therefore, in using roGFP2-ORP1, it is possible that other hydroperoxides could be detected *in vivo* instead of, or alongside H_2_O_2_. This is because ORP1/GPX3 is one of a large class of monomeric GPXs which include those containing either a single redox active selenocysteine (e.g. *Hs*GPX4) or a redox active cysteine pair, the peroxidatic and resolving cysteines, at their active sites (Toppo et al 2009). Collectively, across the monomeric GPXs, the same wide peroxide substrate specificity is encountered and extends to (phospholipid) fatty acid peroxides (Antunes et al 1995; Herbette et al 2002; Flohé et al 2003; Maiorino et al 2007; Toppo et al 2009; Yang et al 2006).

A second class of GPX enzymes, the so-called “classical” GPX enzymes (Avery et al 2004; Toppo et al 2009) (e.g. *Hs*GPX1) are generally selenocysteine-containing, tetrameric forms of the enzyme and display again a wide peroxide substrate specificity but with the exclusion of phospholipid hydroperoxides (Nakamura et al 1974; Madipati and Marnett 1987; Ursini et al 1995; Toppo et al 2009). Of particular relevance to this study, the addition of a putative subunit interaction sequence from classical GPXs can convert ORP1/GPX3 into a “classical” GPX and narrow its substrate specificity to exclude binding to phosphatidylcholine hydroperoxide (Avery et al 2004). Our aim was to construct roGFP2-GPX redox relay sensors that would enable the determination of changes in the levels of both H_2_O_2_ and lipid peroxides (LOOH) of which the most likely class may be phospholipid fatty acid peroxides in the cytosol, nucleus and chloroplasts of plant cells. We achieved this by constructing a modified version of roGFP2-ORP1, which we named roGFP2-synORP1. By using both probes in comparative imaging experiments, we have shown that the intracellular distribution of both peroxide types can be distinguished. This has important implications for the way in which signaling involves LOOH or H_2_O_2_ as both message initiators and transducers. For these peroxides, the potential for their associated signaling pathways to be spatially distinct provides an additional dimension to signaling specificity.

## RESULTS

### Structural considerations

To initiate our study, the amino acid sequences in the subunit interaction region of *Hs*GPX1, *Hs*GPX4 and GPX3/ORP1 were aligned (Fig. 1A; Avery et al 2004) along with the nearest plant *Hs*GPX4-like enzyme reported, *Rs*GPX (Yang et al., 2006; Fig. 1A). The sequence alignments showed two characteristic sequence gaps at similar positions in *Hs*GPX4, *Rs*GPX and ORP1 compared with *Hs*GPX1 (Fig. 1A). A three-dimensional structural alignment between *Hs*GPX1 and *Hs*GPX4 confirmed a disruption of surfaces involved in oligomerization in tetrameric GPXs (Ursini et al., 1995; Fig. 1B). The introduction of these two sequences at the same position in a modified ORP1 greatly diminished its catalysis of phosphatidylcholine hydroperoxide reduction (Avery et al 2004). Therefore, these alignments were used to design a synthetic version of ORP1, hereafter called synORP1 (Fig. 1A).

Experimentally determined protein structures were obtained from the Protein Data Bank (PDB) to evaluate similarities and differences between the electrostatic surfaces of our selected GPXs (Fig. 1C; Supplemental Figure 1). The surface electrostatic properties of the whole protein are important for substrate specificity (Toppo et al 2008; Cozza et al 2017; Gellert et al 2019). Unlike *Hs*GPX1 (Supplemental Figure 1), both *Hs*GPX4 (Supplemental Figure 1) and ORP1 (Fig. 1C) show a large cationic area on their surfaces around their catalytic center which would generate a strong positively charged electrostatic field. *Rs*GPX is structurally like *Hs*GPX4 and has a surface cationic region, although this is not as extensive as in *Hs*GPX4 and ORP1 (Supplemental Figure 1; Fig. 1C). *Hs*GPX1, like synORP1 does not show the large cationic area because it is disrupted by two peptide insertions (Fig. 1A, C; Supplemental Figure 1). From the structural and electrostatic field characteristics (Fig. 1C; Supplemental Figure 1), it was reasoned that both ORP1 / synORP1 and *Hs*GPX1 / *Hs*GPX4 could be pair-moieties to construct redox relay biosensors based on roGFP2 that would display differential hydroperoxide specificities. However, *Hs*GPX4 and *Hs*GPX1 both have single redox-active selenocysteines at their active sites (Maiorino et al 2007; Cozza et al 2017). Since plants do not have selenocysteine-containing proteins (Novoselov et al 2002), and substitution for selenocysteine with cysteine in modified GPXs results in three orders of magnitude less catalytic activity (Maiorino et al 2007; Toppo et al 2009), these were excluded from further analysis.

### Both ORP1 isoforms can oxidize roGFP2

Based on the previous observation that the close proximity of roGFP2 fused with thiol peroxidases can mediate its H_2_O_2_-dependent oxidation (Fig. 2A; Gutscher et al 2009; Morgan et al 2016), *Rs*GPX, ORP1 and synORP1 were separately tethered to the C terminus of roGFP2 using a 32-amino acid linker peptide based on GGSGG repetitions (Gutscher et al 2009). As a control, roGFP2 alone (i.e., unfused to a GPX moiety) was used to monitor the susceptibility of roGFP2 to oxidation *in vitro* in the presence of the peroxides. RoGFP2-ORP1, roGFP2-synORP1, roGFP2-*Rs*GPX and roGFP2 recombinant proteins were over-expressed in and purified from *Escherichia coli* (see Methods; Supplemental Figure 2A). The roGFP2-fusion proteins became much more oxidized in response to H_2_O_2_ and TBP (Fig. 2C, D; Supplemental Figure 3A-D; Supplemental Figure 4A) than unfused roGFP2 control (Fig. 2B; Supplemental Figure 4C). The fusions with the synORP1 moiety did not alter the dual excitation property of roGFP2, which displayed an increased fluorescence peaking at 512 nm from an excitation maximum at 395 nm and a corresponding decrease from the emission maximum at 488 nm (Supplemental Figure 2B). These results demonstrated that the structurally modified synORP1 *in vitro* efficiently catalyzes H_2_O_2_- and TBP-dependent reduction, albeit with a diminished maximum oxidation/reduction ratio compared with roGFP2-ORP1 (Fig. 2E, F): Fully reduced roGFP2-ORP1 showed a 5.3-fold maximum oxidation by both H_2_O_2_ and TBP, while roGFP2-synORP1 was 2.8-fold and 2.6-fold maximally oxidized by H_2_O_2_ and TBP respectively.

**Figure 2.**
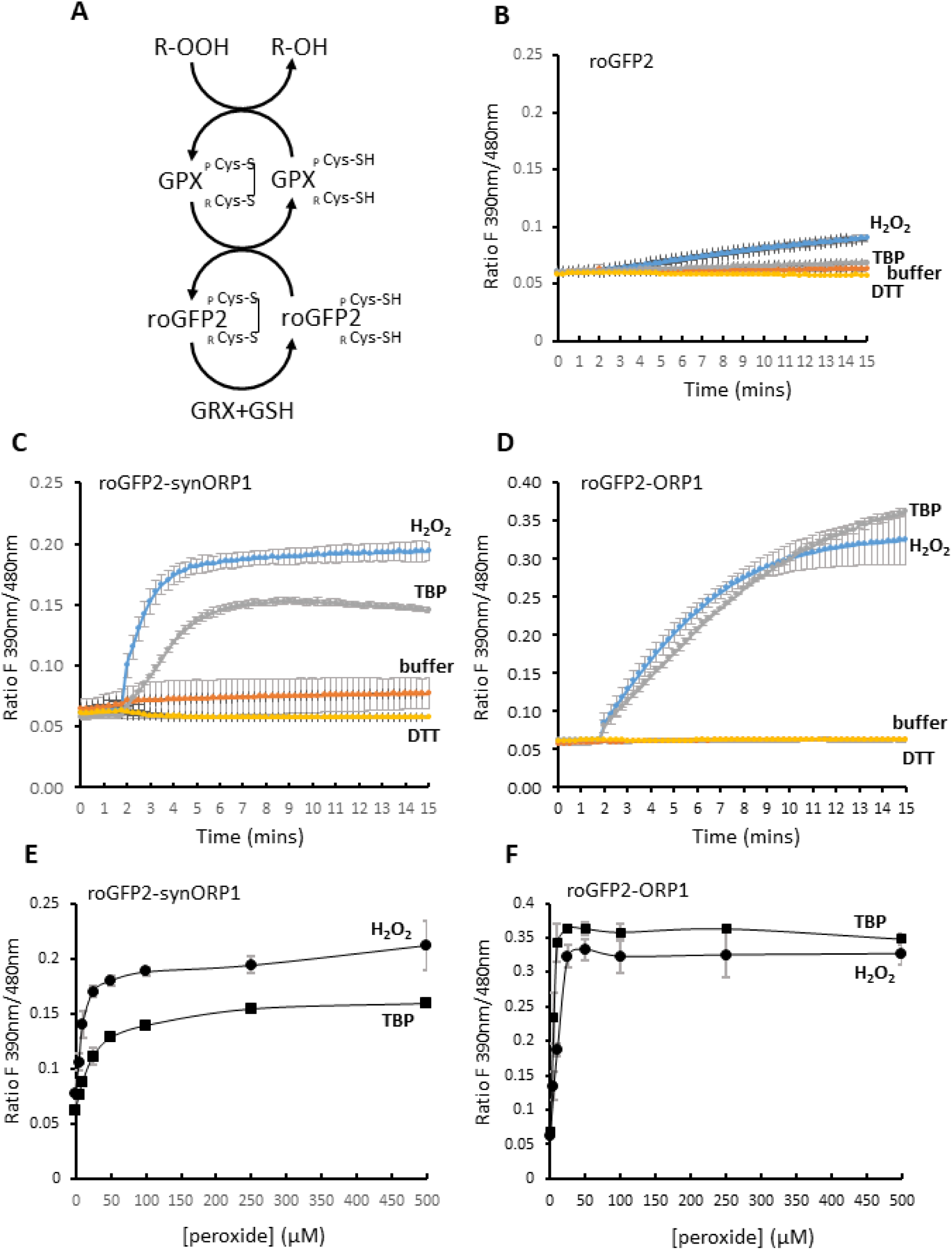
ORP1 isoforms mediate the *in vitro* oxidation of roGFP2 by H_2_O_2_ and TBP. (A) Diagram of the flux of electrons to inorganic or organic hydroperoxides (ROOH) from reduced roGFP2 via GPX in a proximity-based redox relay between redox-active cysteines. *In vitro* oxidation of 5 μM **(B)** roGFP2 by 500 μM H_2_O_2_ or TBP and oxidation of **(C)** roGFP2-synORP1 and **(D)** roGFP2-ORP1 by 250 μM H_2_O_2_ or TBP. Each biosensor was also exposed to 1 mM DTT or buffer alone. Additions of peroxide or DTT was carried out 90 seconds after initiating measurements. Readings for roGFP2-synORP1 and roGFP2-ORP1 are means (± SD) combined from 4-6 technical replicates from two separate preparations. The roGFP2 alone readings are from 3 technical replicates and a single preparation. Maximum oxidation (ratios of fluorescence emissions at 530 (± 20) nm from excitations at 390 (± 10) nm and 480 (± 10) nm) attained for (**E**) roGFP2-synORP1 and (**F**) ro-GFP2-ORP1 after exposure to H_2_O_2_ or TBP at final concentrations of 0 (control), 5 μM, 10 μM, 25 μM, 50 μM, 100 μM, 250 μM and 500 μM. Time course plots for each treatment can be viewed in Supplemental Figure 3. The degree of replication was the same as in the legends of B-D.

The particularly wide substrate specificity of *Rs*GPX (Yang et al 2006) was considered suitable as the basis for making an additional control of a disabled peroxidase moiety tethered to roGFP2. This was intended to be a generic control that would prove suitable for both roGFP2-ORP1 and roGFP2-synORP1 comparisons *in vivo* (see below). A mutant was made in which the *Rs*GPX moiety active site peroxidatic cysteine was changed to a serine (C71S). This mutation abolished the *in vitro* enzyme activity of the GPX moiety fused to roGFP2 (Supplemental Figure 4B).

### Response of the biosensors to *in vitro* liposome oxidation by soybean lipoxygenase

To test the biosensors’ response towards phospholipid fatty acid peroxide substrates, we evaluated roGFP2-ORP1, roGFP2-synORP1 and roGFP2-*Rs*GPX oxidation during the lipoxygenase (LOX)- catalyzed oxidation of unilamellar liposomes prepared from a mixture of 65-75% soybean phospholipids comprising ∼20% L-α-Phosphatidylcholine; ∼14% L-α-Phosphatidylethanolamine and ∼20% Inositol phosphatides (Scholfield 1981; Tayeb et al 2017; Fig. 3A; see Methods). The LOX-dependent production of LOOH was monitored optically by the appearance of lipid-based conjugated dienes (Fig. 3B; Reeder et al 2004). In a mix of liposomes and reduced roGFP2-biosensors, the addition of LOX initiated roGFP2 oxidation (Fig. 3C). Notably, a large increase in the rate of roGFP2-ORP1 and roGFP2-*Rs*GPX oxidation resulting in a doubling of their oxidation state was measured compared with a maximum increased oxidation of 20% of starting values for roGFP2-synORP1, but only a negligible increase in oxidation of the roGFP2 control was observed (Fig. 3C). Therefore, from these *in vitro* trials, we confirmed that roGFP2-ORP1 and roGFP2-synORP1 tested as a pair could be used to discriminate between LOOH and H_2_O_2_.

**Figure 3.**
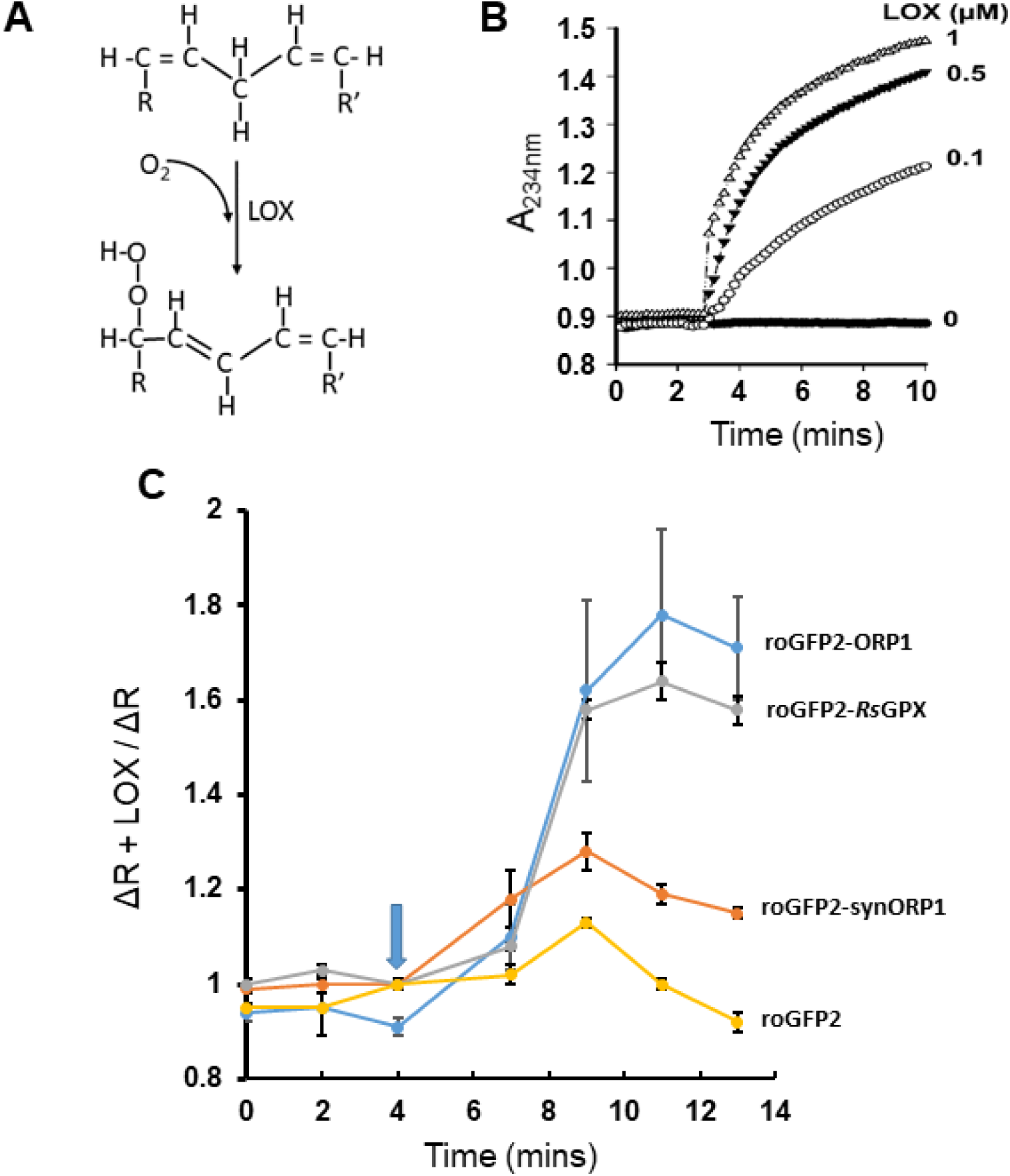
ORP1 but not synORP1 mediates the *in vitro* oxidation of roGFP2 by PFA-OOH in unilamellar liposomes. (A) The formation of lipid peroxide catalyzed by the dioxygenase reaction of LOX. R and R’ represent the carbon chains of the rest of the lipid molecule (after Tayeb et al 2017). **(B)** Increase in a LOX-dependent manner of absorbance at 234 nm caused by the generation of conjugated dienes during incubation of 0, 0.1 µM, 0.5 µM and 1µM soybean LOX with 200 μg ml^-1^ liposomes. **(C)** Response of recombinant roGFP2 probes (5µM) as indicated in response to LOX (1µM)-catalyzed liposome oxidation. The arrow indicates when the enzyme was added. Data are means of three sets of measurements each from a separate liposome preparation. Each reaction was run in parallel with a control in which no LOX was added. Data were normalized for each pair to the starting time zero value and expressed as the ratio ΔR **+** LOX **/** ΔR (control) for each time point. Values shown are the means (± SD; n=3).

### The roGFP2-ORP1 and -synORP1 probes function *in vivo*

To confirm that the roGFP2-ORP1, roGFP2-synORP1 can respond at the subcellular level *in vivo*, we examined their fluorescence properties under extreme oxidation and reduction conditions in chloroplasts, cytosol and nuclei of *Nicotiana benthamiana* abaxial epidermal cells (Supplemental Figure 5A, B). This was done by following transient expression of the constructs between 3 and 4 days after agro-inoculation of leaves and then imaging leaf discs under a confocal scanning laser microscope (CSLM; Exposito-Rodriguez et al 2017; see Methods).

Under extreme oxidizing conditions, these redox relay probes could be subject to direct alterations in the redox state of the roGFP2 moiety irrespective of their tethered peroxidase. Independent oxidation of the roGFP2 moiety has been raised as a potentially confounding problem of data interpretation (Schwarzlander et al 2016). This notion was tested *in vivo* using roGFP2-*Rs*GPXC71S. To achieve an extreme oxidation and reduction of the biosensors *in vivo*, agroinfected leaf discs were infiltrated with 100 mM H_2_O_2_, 20 mM dithiothreitol (DTT) or water (control) (see Methods). The F405/F488 530nm emission ratio (see Introduction) increased in all three compartments for all biosensors when oxidized by this high concentration of H_2_O_2_ (Supplemental Figure 5A, B). Imposition of reducing conditions by DTT decreased the ratio to the same low value in all the compartments (Supplemental Figure 5A, B). It was clear from these initial experiments under extreme conditions that the probes’ roGFP2 moiety can, under certain conditions, become fully oxidized independent of their tethered GPX. This illustrates that any generalized impacts of a particular stress or treatment on cellular redox state can be discounted using a control such as roGFP2-*Rs*GPXC71S and consequently, the (syn)ORP1-based probes’ oxidation be attributed to *in vivo* changes in peroxide levels.

### Differential biosensor responses to the accumulation of H_2_O_2_ and LOOH caused by photoinhibition

The herbicide 3-(3’,4’-dichlorophenyl)1,1-dimethylurea (DCMU) binds specifically to the Quinone B (Q_B_) site of the D1 protein of Photosystem II (PSII) reaction centers, thereby preventing electron transfer from Q_A_ to Q_B_ (Bowyer et al 1991). In the light, binding of DCMU to the PSII reaction center results in the formation of the radical pair P+680 Pheophytin^-^ whose recombination produces the excited triplet state of P680 (P680^3^), which reacts with O_2_ to form ^1^O_2_ (Rutherford and Krieger-Liszkay 2001; Flors et al 2006; Dogra and Kim 2020). Consequently, the increased ^1^O_2_ can give rise to LOOH, principally as (phospholipid) fatty acid hydroperoxides (Triantaphylidès et al 2008), which are substrates for GPX3/ORP1 (Avery et al 2004; Fig. 3C). Therefore, it was reasoned that when using the roGFP2-synORP1 and roGFP2-ORP1 in parallel, comparative fluorescence data would allow a discrimination between H_2_O_2_ and LOOH levels *in vivo*. In a pilot experiment using only cytosol-targeted roGFP2-ORP1, treatment of transiently expressing leaf discs with DCMU followed by exposure to a high light intensity (HL; see Methods) caused oxidation of the probe after 2.5h (Fig.4A) and was associated with photoinhibition, as determined by chlorophyll fluorescence (Fig.4B). In subsequent experiments a 3h HL exposure was carried out in the presence or absence of DCMU (Fig. 5).

**Figure 4.**
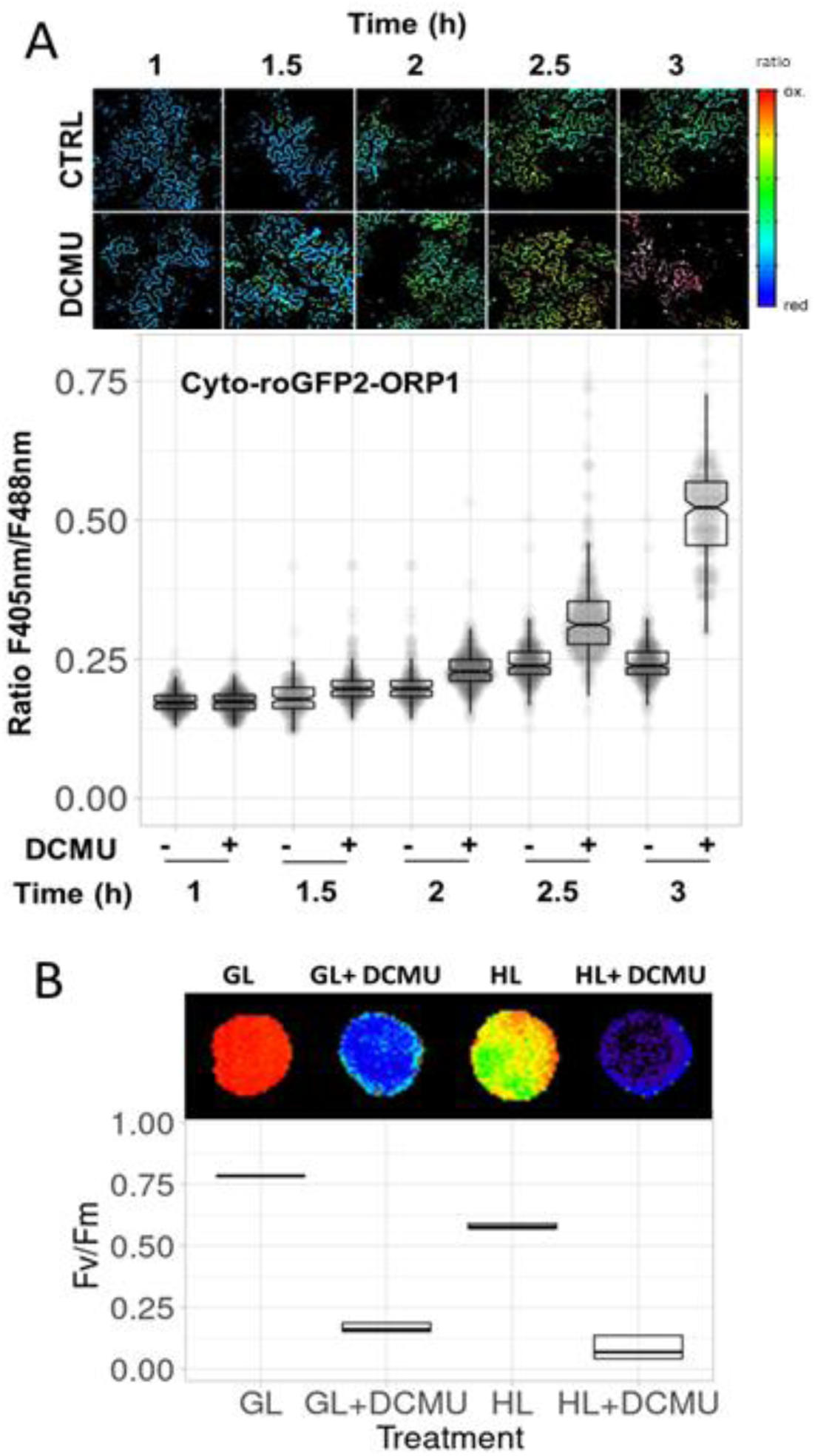
The timing of the *in vivo* response of roGFP2-ORP1 to DCMU and HL treatment. Treatment of leaf discs of *N. benthamiana* transiently expressing cytosol-targeted roGFP2-ORP1 with 10µM DCMU for 10 min in the dark followed by exposure to HL (1000 µmol m^-2^ s^-1^; +) compared with HL alone (-). (**A**) Oxidation of the probe was monitored in abaxial epidermal cells at the indicated times as a rise in the F405 nm/488 nm ratio. The plots are compiled from ≥ 150 readings per sample. (**B**) Photoinhibition as determined by chlorophyll fluorescence imaging using the Fv/Fm CF parameter for maximum PSII quantum efficiency (Exposito-Rodriguez et al 2017) in leaf discs exposed to growth light (GL) or HL ± DCMU for 3h.

**Figure 5.**
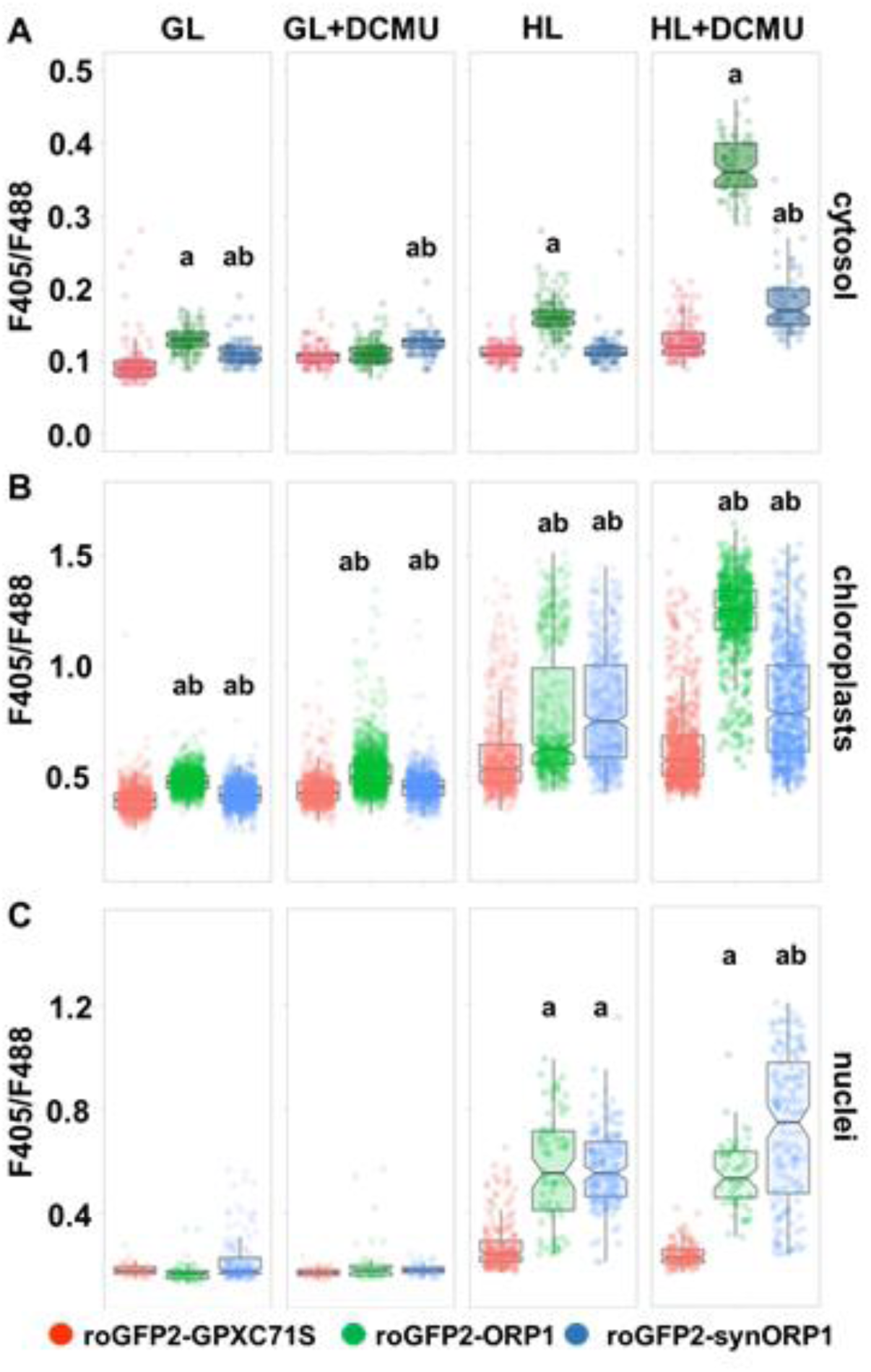
Monitoring the oxidation state of the biosensors in response to herbicide-induced inhibition of photosynthesis. *Nicotiana benthamiana* leaf discs expressing the indicated roGFP2 biosensor construct in cytosol (**A**), chloroplast stroma (**B**) and nuclei (**C**) were placed in buffer or buffer supplemented with 10 µM DCMU to inhibit photosynthetic linear electron flux. After 1 h, the samples were exposed to 3 hours of growth light (GL) or high light (HL; see Methods). The number of acquired image data points collected per treatment for each subcellular compartment are given in Supplemental Table 1, but for cytosol were n ≥ 113; chloroplasts, n ≥ 590; nuclei, n ≥ 54. Data are given as jitter dots and are from 4 discs per condition and 3 independent experiments. Boxplots reflect the data distribution, and a horizontal line indicates the median. Vertical line indicates for each median the 95% confidence, which is calculated from 1000 bootstrap samples. The p-values were determined by a randomization test; ‘a’ represents significant differences between roGFP2-GPXC71S (control) and roGFP2-ORP1 and roGFP2-synORP1 (p-values < 0.001) and ‘b’ differences between roGFP2-ORP1 (control) and roGFP2-synORP1 (p-values <0.001).

When the probes were targeted to the cytosol, HL exposure combined with DCMU treatment promoted oxidation of roGFP2-ORP1, but much less oxidation of roGFP2-synORP1 compared with roGFP2-*Rs*GPXC71S (P <0.001; Fig. 5A). It was inferred that LOOH strongly accumulated in this compartment under conditions of severe photoinhibition to a much greater degree than any increase in H_2_O_2_ levels. In the absence of DCMU, cells exposed to 3h HL only, showed a significant (p<0.001) oxidation of roGFP2-ORP1, but no increased oxidation of roGFP2-synORP1 (Fig. 5A), meaning that under more moderate photoinhibitory conditions, cytosol showed increased LOOH levels but H_2_O_2_ levels were unaffected.

Likewise, in chloroplasts exposed to HL + DCMU, there was greater oxidation of roGFP2-ORP1 compared with roGFP2-synORP1 (Fig. 5B) and again was largely attributed to increased LOOH levels and to a smaller increase in H_2_O_2_ levels. In abaxial cells exposed to HL alone, the chloroplast-located probes showed the same degree of increased oxidation (Fig. 5B), which was consistent with increased levels of H_2_O_2_ and LOOH in this compartment under these conditions. In nuclei, the oxidation of roGFP2-synORP1 in DCMU + HL-treated cells was significantly (p<0.001) greater than roGFP2-ORP1 (Fig. 5C) indicating that there was preferential accumulation of H_2_O_2_ in nuclei under these conditions and it was further concluded that nuclei did not accumulate LOOH irrespective of the degree of photoinhibition.

## DISCUSSION

ORP1/GPX3, whether tethered to roGFP2 (Gutscher et al 2009) or in its native form (Delaunay et al 2002; Avery et al 2004), catalyzes the reduction of H_2_O_2_, synthetic organic peroxides (e.g. TBP) and lipid peroxides, the most likely being (phospholipid) fatty acid hydroperoxides such as phosphatidylcholine hydroperoxide (Figs. 2, 3; Supplemental Figures 3, 4). In its native form, ORP1 is monomeric (Avery et al 2004) and this exposes a surface area of cationic charge (Fig. 1C). This is important in allowing binding of negatively charged phosphate in the polar head group of phospholipid fatty acid peroxides, in turn, facilitating the interaction of the peroxide(s) in the aliphatic chain with the enzyme’s catalytic site (Avery et al 2004; Toppo et al 2009; Cozza et al 2017). Being tethered to roGFP2 does not hinder this interaction (Fig. 3C). Direct access by H_2_O_2_ and TBP to the catalytic triad does not depend upon a cationic surface charge since its disruption by a α-helix and loop sequence (Fig. 1A; Avery et al 2004) in synORP1 does not substantially inhibit reduction of these peroxides, but does drastically diminish the reduction of (phospholipid) fatty acid hydroperoxides in unilamellar liposomes (Fig 3C). In addition, the synORP1 moiety of this probe may exist in a dimeric form, which may further obscure the surfaces that interface with LOOH (Avery et al 2004). We have not determined if synORP1-roGFP2 is a dimer but the formation of multimers for the redox relay H_2_O_2_ biosensor (ro)GFP2-TsaΔC_R_ does not hinder its function (Morgan et al 2016).

Based on its peroxide affinities (Figs. 2, 3), roGFP2-synORP1 can be used to determine *in vivo* changes in H_2_O_2_ with more certainty than reliance solely on roGFP2-ORP1. The increased oxidation state of roGFP2-synORP1 in abaxial cell chloroplasts subjected to 3h HL (Fig. 4B) is consistent with HL-induced production of H_2_O_2_ identified in isolated chloroplasts determined using the H_2_O_2_ specific dye Amplex Red (Mubarakshina et al 2010) and in HL-exposed epidermal pavement cells from *N. benthamiana* and *Arabidopsis thaliana* using variants of HyPer (Exposito-Rodriguez et al 2017; Ugalde et al 2021a, b).

However, a more valuable strategy is to compare the behavior of the two probes in parallel (Fig. 5). It is possible to infer changes in the steady state levels of both H_2_O_2_ and LOOH, most likely (phospholipid) fatty acid hydroperoxides, in stressed cells depending on the probes’ differential oxidation in these conditions. Both roGFP2-synORP1 and roGFP2-ORP1 were oxidized to the same degree in the HL-exposed chloroplasts (Fig. 5B), indicating that only significant increased amounts of H_2_O_2_, compared with GL conditions, were present. In contrast, the more severe photo-oxidative stress induced by HL + DCMU treatment produced a much higher degree of oxidation of roGFP2-ORP1 than roGPFP2-synORP1 in the chloroplast stroma (Fig. 5B). This could be attributed to increased levels of LOOH. Short incubation times (≤ 1h) of HL+DCMU lead to suppression of H_2_O_2_ levels attributed to the inhibition of electron transport-driven Mehler reaction (Mubarakshina et al 2010; Exposito-Rodriguez et al 2017; Ugalde et al 2021a). In contrast, the 3h incubation time used to drive increased ^1^O_2_ production did cause an increase in roGFP2-synORP1 oxidation but not to the same extent as the oxidation of roGFP2-ORP1 (Fig. 5B). Therefore, under these conditions of prolonged DCMU-induced photo-oxidative stress, the stromal steady state H_2_O_2_ levels do rise. The source of this H_2_O_2_ is not clear, but may be due to interconversions among ROS driven from ^1^O_2_ production in the presence of antioxidants such as ascorbate (Kramarenko et al 2006; Lelieveld et al 2021).

Significantly greater oxidation of roGFP2-ORP1 than roGFP2-synORP1 was evident in the cytosol of leaf tissue exposed to HL+DCMU (Fig. 5A) indicating that LOOH was the predominant peroxide class that accumulated in this compartment. Under oxidative stress, lipid peroxidation arises from the plasma membrane (Cozza et al 2017, de Dios Alché 2019), although how HL+DCMU would influence these processes is not clear. The export of fatty acid peroxides, reactive electrophile species and oxylipins from chloroplasts suffering severe photooxidative stress (Wasternack and Feussner 2018; Munoz and Munne-Bosch, 2020) may have initiated further lipid peroxidation in the cytosol as can peroxynitrite (de Dios Alché 2019). As a note of caution, we cannot rule out that the biosensors were oxidized by peroxynitrite (Schwarzlander et al 2016; Muller et al 2017), although the differential patterns of expression observed (Fig. 5A, C) strongly suggest that this reactive nitrogen species is not prevalent in our experimental system.

If the source of lipid peroxides detected in the cytosol (Fig. 5A) comes from severely photo-inhibited chloroplasts (Fig. 4B), then to our knowledge there is no study pointing to how fatty acid peroxides, the presumed breakdown products of (phospholipid) fatty acid hydroperoxides formed in membranes, could be exported from chloroplasts. We speculate that that fatty acid hydroperoxides could perhaps use the same membrane transporters as for unmodified fatty acids (Li et al 2016). Under oxidative stress (e.g. produced by long dark periods and carbon starvation) the transport of fatty acids from chloroplasts markedly increases (Fan et al 2017). In support of this suggestion, transcriptome studies in Arabidopsis that unite the various means of eliciting ^1^O_2_-induced oxidative stress show many differentially expressed genes encoding ABC transporters (Willems et al 2016; Bode et al 2016; Koh et al 2022). An additional possibility is that lipid peroxides could be conjugated to or reduced by glutathione (as GSH) by a multiplicity of Glutathione-S-Transferase (GST) isoforms (Dixon and Edwards 2010; de Dios Alché 2019). Many GST genes are highly expressed under the same oxidative stress conditions (Willems et al 2016; Koh et al 2022). However, notwithstanding how the accumulation of LOOH occurs in the cytosol, their presence likely indicates an advanced state of oxidative stress that would initiate programmed cell death (Mullineaux and Baker 2010; Willems et al 2016; Wasternak and Feussner 2018; Koh et al 2022).

The preferential oxidation of roGFP2-synORP1 compared with roGFP2-ORP1 in nuclei of epidermal pavement cells exposed to HL or HL+DCMU (Fig. 5C) suggests again a capacity for nuclei under stress to make H_2_O_2_ or for it to be transferred from chloroplasts or elsewhere. The transfer of H_2_O_2_ from photosynthetically active chloroplasts in close physical association with the nuclei in HL conditions was an important conclusion from a more comprehensive analysis conducted using the H_2_O_2_ biosensor HyPer (Exposito-Rodriguez et al 2017; Mullineaux et al 2020; Breeze and Mullineaux 2022). However as stated above, the 3h DCMU treatment in in this study (Fig. 5A-C) was associated with increased oxidation of nuclear-located roGFP2-synORP1 and was most likely caused by increased H_2_O_2_ production in photo-inhibited chloroplasts and its transfer to the adjacent nucleus (Fig. 4A; Exposito-Rodriguez et al 2017; Mullineaux et al 2020; Breeze et al 2022). The passage of H_2_O_2_ between compartments and has been proposed, based on physico-chemical considerations, to be facilitated by water channel aquaporins (Wang et al 2020). This proposal has been experimentally confirmed in plant suspension culture cells in an elegantly designed experiment (Ahmed et al 2023).and thus providing support for the notion that H_2_O_2_ can be a transducing signal between closely associated chloroplasts and the nucleus.

## METHODS

### Protein structures, molecular modelling, and electrostatic calculations

MEGA6 was used for the alignment of amino acid sequences retrieved from NCBI (Tamura et al 2013). Structures for *Hs*GPX1 (2F8A), *Hs*GPX4 (2OBI) and *Sc*GPX3/ORP1 (3CMI) were obtained from the Protein Data Bank (https://www.rcsb.org). The *Rs*GPX and synORP1 3D structures were calculated by homology modelling using ProMod3 (Studer et al 2018) implemented in the Swiss Model server using PDB entry 2v5y as template (https://swissmodel.expasy.org; Schwede 2017).

The electrostatic properties of the proteins were computed from the pdb files. The reconstruction of any missing atoms, the addition of hydrogens, the assignment of atomic charges and radii were performed using pdb2pqr with the Amber force field (Dolinsky et al 2007). The electrostatic parameters were calculated using the Adaptive Poisson-Boltzmann Solver (APBS) (Baker et al 2001) within the vmd (visual molecular dynamics) software package (Humphrey et al 1996) implemented in PyMOL (PyMOL:Molecular Graphics System, Version 2.1.1 Schrödinger, LLC). The following parameters were used: 150 mM mobile ions, solvent dielectric constant: 78.54, temperature: 298.15 K. Images were rendered depicting the secondary structures of the proteins, the electrostatic potential mapped to the surface of the proteins (from -2 in red to 2 KT·e^−1^ in blue).

### Plasmids, DNA Constructs

All synthetic DNA fragments were purchased from GeneWiz (Takeley, Essex, UK). roGFP2-ORP1, roGFP2-synORP1, GPXC71S-roGFP2 and unfused roGFP2 (Supplemental Data 1) were cloned into the *Nde*I and *Hind*III sites in the pET28a(+) expression vector (Novagen, Cambridge, UK) which resulted in a N-terminal fusion with a polyhistidine tag (Supplemental Data 1). For transient expression in *Nicotiana benthamiana*, the sequences described above were cloned in frame with the signal peptide NLS (in plasmid pICH86988; Addgene (www.addgene.org) # 48076) and a transit peptide (in plasmid pICH78133; Addgene # 50292) to target the biosensor to the nuclei and chloroplast stroma respectively. MoClo Golden Gate cloning (Weber et al 2011) was used to assemble DNA fragments in pICH86988, which is a binary Ti vector) using AATG-GCTT sequences between the CaMV 35S promoter and *Agrobacterium tumefaciens* octopine synthase terminator *via Bsa*I restriction enzyme sites.

All Ti plasmids harboring the roGFP2-based probes can be found in the Addgene plasmid depository (www.addgene.org) with the following ID codes: 191677 (pICH-roGFP2-ORP1), 191680 (pICH-roGFP2-synORP1), 191698 (pICH-roGFP2-GPXC71S), 191699 (pICH-chl-roGFP2-ORP1), 191701 (pICH-chl-roGFP2-synORP1), 191703 (pICH-chl-roGFP2-GPXC71S), 191758 (pICH-nuc-roGFP2-ORP1), 191759 (pICH-nuc-roGFP2-synORP1) and 191760 (pICH-nuc-roGFP2-GPXC71S).

### Isolation of Recombinant Proteins

After the pET28a-roGFP2-GPX fusion expression plasmids were transformed into *Escherichia coli* BL21 (DE3) (Fisher Scientific, Loughborough, UK), a pre-culture of 5 mL volume was grown at 37 °C overnight. This pre-culture was added to 500 mL Luria Bertani medium and grown to an OD_600_ of 0.6–0.8. To avoid misfolding, the temperature was subsequently decreased to 18 °C and 0.2 mM isopropyl-thio-β-D-galactoside (IPTG) was added for overnight induction. For recombinant target protein purification, the cell pellet derived from the culture was resuspended in a column binding buffer (50 mM Tris, 500 mM NaCl, pH 8, at 10 mL/g of cell pellet) including protease inhibitors (complete protease inhibitor cocktail tablet, Roche Life Sciences, UK). The suspension was lysed using an Emulsiflex extruding homogenizer (Avestin, Ottawa, Canada) and cell debris removed by centrifugation at 4 °C, 10,000 xg for 30 min. The supernatant was further clarified by ultra-centrifugation (45,000 xg for 1 h) and passed through a 0.45 µm pore filter. Nickel affinity chromatography columns (His-Trap HP columns, GE Healthcare, Chalfont St Giles, Bucks., UK) were used to purify the polyhistidine-tagged roGFP2-GPX fusion proteins with a gradient elution (50 mM Tris, 500 mM NaCl, from 10 mM to 500 mM imidazole, pH 8) using the liquid chromatography system ÄKTA (Amersham Biosciences, Little Chalfont, Bucks., UK).

After desalting by molecular weight cut-off dialysis (Pierce Slide-A-Lyzer, Fisher Scientific), the proteins were stored at −80 °C. Protein concentrations were quantified using the Pierce BCA Protein Assay Kit (Fisher Scientific). Protein expression and purity were examined using SDS-PAGE (Supplementary Fig. 4).

### *In Vitro* Fluorescence-based Assays

Recombinant proteins were diluted into standard reaction buffer (100 mM potassium phosphate, 5 mM EDTA, pH 7.0, degassed and then saturated with N_2_) to a final concentration of 5 μM in 95µl aliquots and immediately frozen on dry ice. Oxidation with H_2_O_2_ or TBP at the concentrations and times indicated in the legends of Fig. 2 and Supplemental Figure 3 and no more than three reactions were run at any one time and their fluorescence determined at 15 sec intervals. The fluorescence emission of roGFP2 (530 nm ± 20 nm) after excitation at 390 (±10) and 480 (±10) nm was measured using a plate reader (either a CLARIOstar+ or a FLUOstar Omega; BMG Labtech). The ratio of the emissions was calculated and plotted against time as described by Aller et al (2017).

### *In vitro* liposome oxidation by soybean lipoxygenase

Liposomes were prepared from soybean lecithin (50 mg; L-α-Phosphatidylcholine; Sigma-Aldrich P5638) as previously described, with minor modifications (Reeder and Wilson, 2005). Briefly, crude multilamellar liposomes were generated from sonication of 5 mg/ml soybean lecithin, using a sonicating water bath, until particulates were no longer visible. Unilamellar liposomes were made by passing the crude multilamellar liposome preparation through an Emulsiflex-C3 extruding homogenizer (Avestin, Ottawa, Canada) fitted with a drain disk (25 mm diameter) and two layers of Whatman nucleopore polycarbonate membranes (25 mm diameter with 100 nm pore size. The liposomes were passed through the membrane a total of ten times to form small unilamellar liposomes of ∼90-120 nm in diameter. The oxidation of liposomes (200 μg ml^-1^) to generate phospholipid hydroperoxides was catalyzed by soybean lipoxygenase (Sigma-Aldrich L7395) in 50 mM sodium phosphate buffer pH 7.4. Oxidation of liposomes was measured by the appearance of lipid-based conjugated dienes, which form concurrent to the phospholipid peroxides (Flors et al 2006), measured optically at 234 nm using a Tecan Infinite M200Pro plate reader.

### Agroinfection of *N benthamiana* leaves and exposure to HL and DCMU

*N. benthamiana* plants were grown, used and inoculated with disarmed *Agrobacterium tumefaciens* GV3850::mp90 harboring the roGFP2 based biosensor constructs in binary Ti plasmids along with the same strain carrying the P19 post-transcriptional gene silencing suppressor, as described previously (Exposito-Rodriguez et al 2017). One cm diameter leaf discs were cut from 3-4 days post-inoculation leaves and floated on 1/2 MS media with or without 10 µM DCMU in 12-well clear multi-well plates. The discs were incubated in the dark for 10 mins and then exposed to either GL (a photosynthetically active photon flux density (PPFD) of 150 µmol m^-2^ s^-1^) or HL (1000 µmol m^-2^ s^-1^ PPFD) for 3h from a white light emitting diode (LED) array (Isolight 4000; Technologica Ltd, Colchester, UK). The intensity of the light and the temperature were measured with a light meter (SKP200, Skye, Powys, UK) and a homemade thermocouple, respectively. The degree of photoinhibition was determined using chlorophyll fluorescence (CF) imaging to determine maximum PSII quantum efficiency as previously described (Exposito-Rodriguez et al 2017).

### Live Cell Confocal Imaging

The procedures and conditions for confocal laser scanning microscopy have been described in detail previously (Exposito-Rodriguez et al 2017). Briefly, for image acquisition, a Nikon A1si confocal microscope was used equipped with a Nikon S Fluor 20x (NA 0.75, WD 1 mm) objective. The intensity ratio image data were acquired using one-way sequential line scans of two temporally separated excitation lines (405nm and 488nm). For biosensors based on roGFP2, the ratio of 405/488-nm laser power was kept constant at 3:1 and emission collected with one detector at 540/530 nm. No offset was used, and pinhole size was set between 1.2 and 2 times the Airy disk size of the used objective, depending on signal strength. Chloroplast stroma localization of the biosensors was confirmed by chloroplast auto-fluorescence with excitation at 640 nm and emission at 670–720 nm.

### Image Processing

Images were analyzed as described by Gutscher et al. (2008). Briefly, images were smoothed using a Gaussian filter, and pixels containing the biosensor signal separated from the background using Otsu’s method (Otsu, 1979). A ratio image was created by dividing the 488 nm image by the 405 nm image pixel by pixel and displayed using false colors.

### Statistical Analysis

Data analysis was done using PlotsOfData (https://huygens.science.uva.nl/PlotsOfData; Postma and Goedhart, 2019). Data were visualized using box plots and a quasi-random distribution of data.

Corresponding p-values were produced using randomization tests (Hooton, 1991; Nuzzo, 2017).

## ACKNOWLEDGEMENTS

The authors acknowledge the support of the UK Biotechnology and Biological Science Research Council (grant BB/P026656/1), the advice of Dr Richard W. Strange during the course of this work and the gifts of plasmids *via* Addgene from Sylvester Marillonet (Leibniz Institute for Plant Biochemistry, Germany) and Nicola Patron (John Innes Centre, UK).

---

**Supplemental Table 1.**
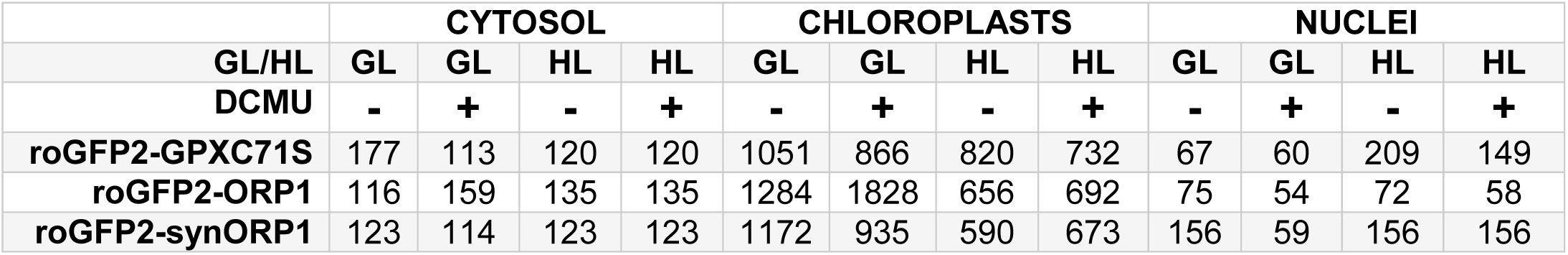
The number (n) of data points from compartment specific ratio images acquired from the four combinations of light intensity and DCMU treatments.

|  | CYTOSOL |  |  |  | CHLOROPLASTS |  |  |  | NUCLEI |  |  |  |
| --- | --- | --- | --- | --- | --- | --- | --- | --- | --- | --- | --- | --- |
| GL/HL | GL | GL | HL | HL | GL | GL | HL | HL | GL | GL | HL | HL |
| DCMU | - | + | - | + | - | + | - | + | - | + | - | + |
| roGFP2-GPXC71S | 177 | 113 | 120 | 120 | 1051 | 866 | 820 | 732 | 67 | 60 | 209 | 149 |
| roGFP2-ORP1 | 116 | 159 | 135 | 135 | 1284 | 1828 | 656 | 692 | 75 | 54 | 72 | 58 |
| roGFP2-synORP1 | 123 | 114 | 123 | 123 | 1172 | 935 | 590 | 673 | 156 | 59 | 156 | 156 |

**Supplemental Figure 1.**
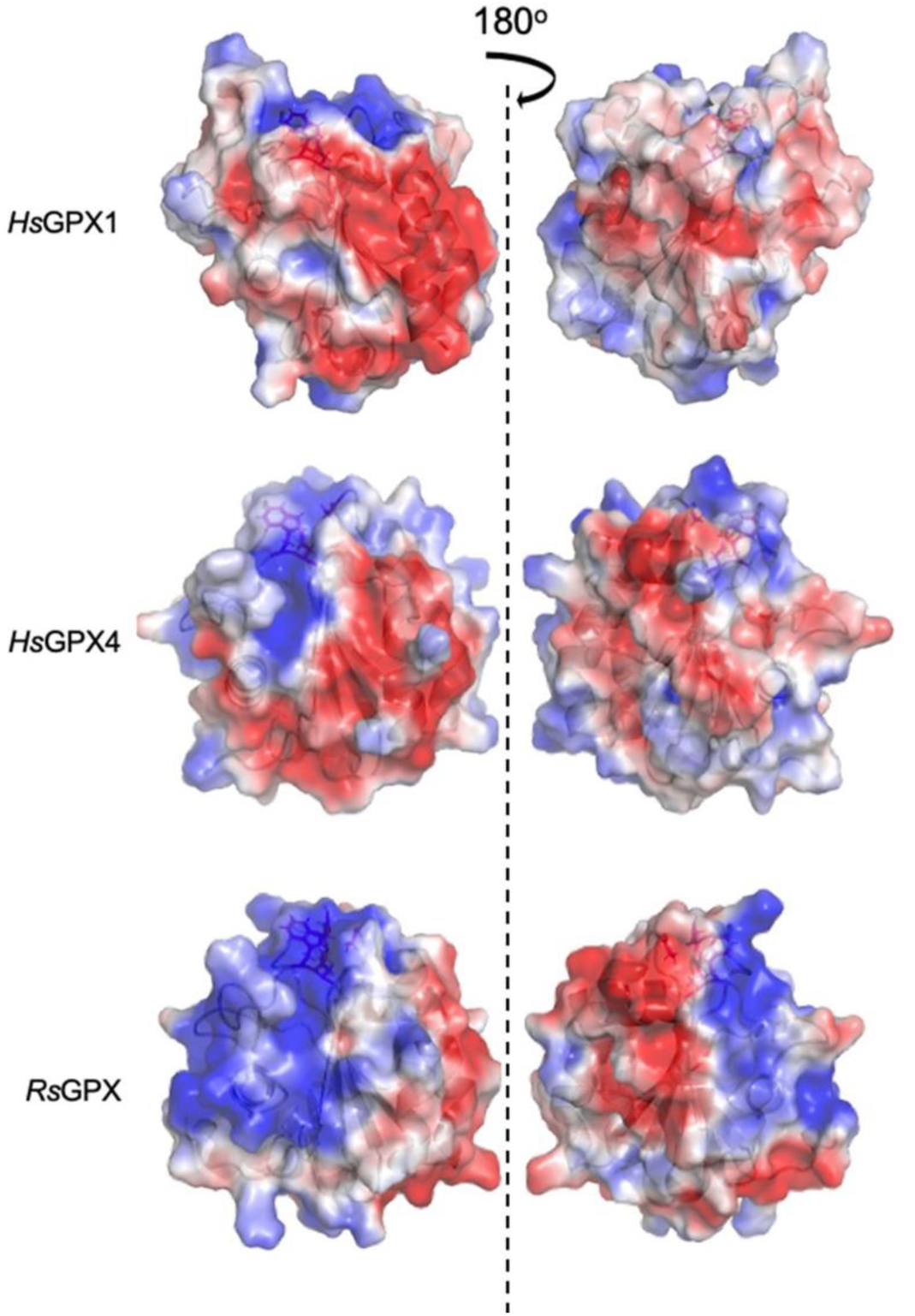
Comparison of the surface electrostatic potentials of *Hs*GPX1, *Hs*GPX4 and *Rs*GPX. Surface electrostatic potentials were calculated using the X-ray crystal structures of *Hs*GPX1 (PDB 2F8A) and HsGPX4 (PDB 2OBI) and simulated by homology modelling of *Rs*GPX using ProMod3 (see Methods). Two representations of each structure are shown rotated by 180 degrees. The electrostatic gradient is shown from the most negatively charged surface (red color;-2 KT·e^−1^) to the most positively charged surface (blue color;2 KT·e^−1^).

**Supplemental Figure 2.**
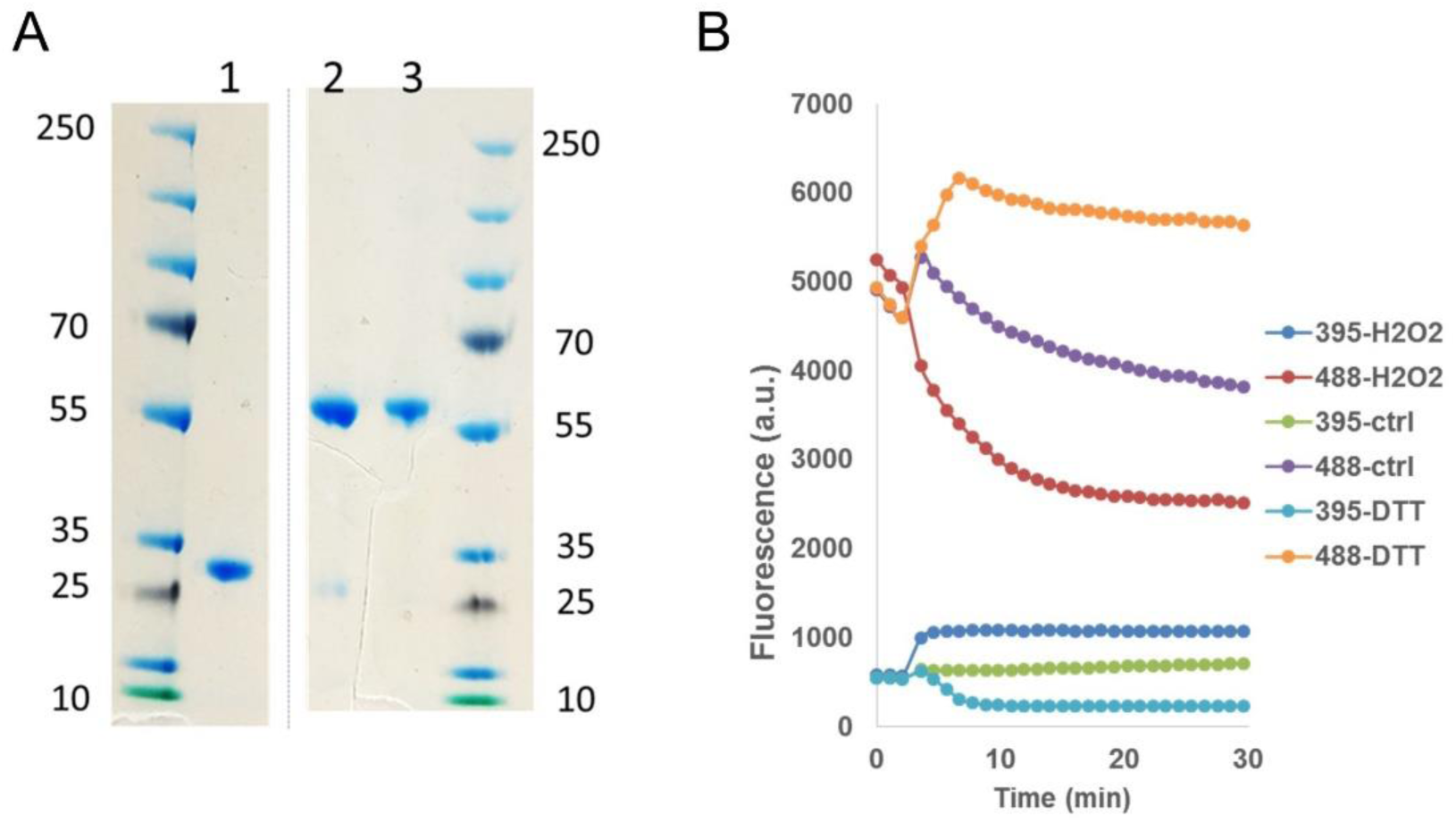
**(A) Confirmation of the purity of the recombinant roGFP2-based biosensor proteins expressed in and purified from *E. coli*.** The indicated recombinant proteins were resuspended in SDS sample buffer, boiled for 5 min and subjected to SDS-PAGE analysis. Lane (1) roGFP2, (2) roGFP2-ORP1 and (3) roGFP2-synORP1. **(B) The synORP1 moiety did not alter the dual excitation property of roGFP2.** 5 μM reduced roGFP2 -synORP1 displayed an increased fluorescence at 512 nm from excitation at 395nm (395-H2O2) and a corresponding decrease from 488nm (488-H2O2) when oxidized with 100μM H_2_O_2_. Treatment with 10 µM DTT produced the converse result (see 488-DTT and 395-DTT). The remaining plots are untreated roGFP2-ORP1 (395-ctrl and 488-ctrl).

**Supplemental Figure 3.**
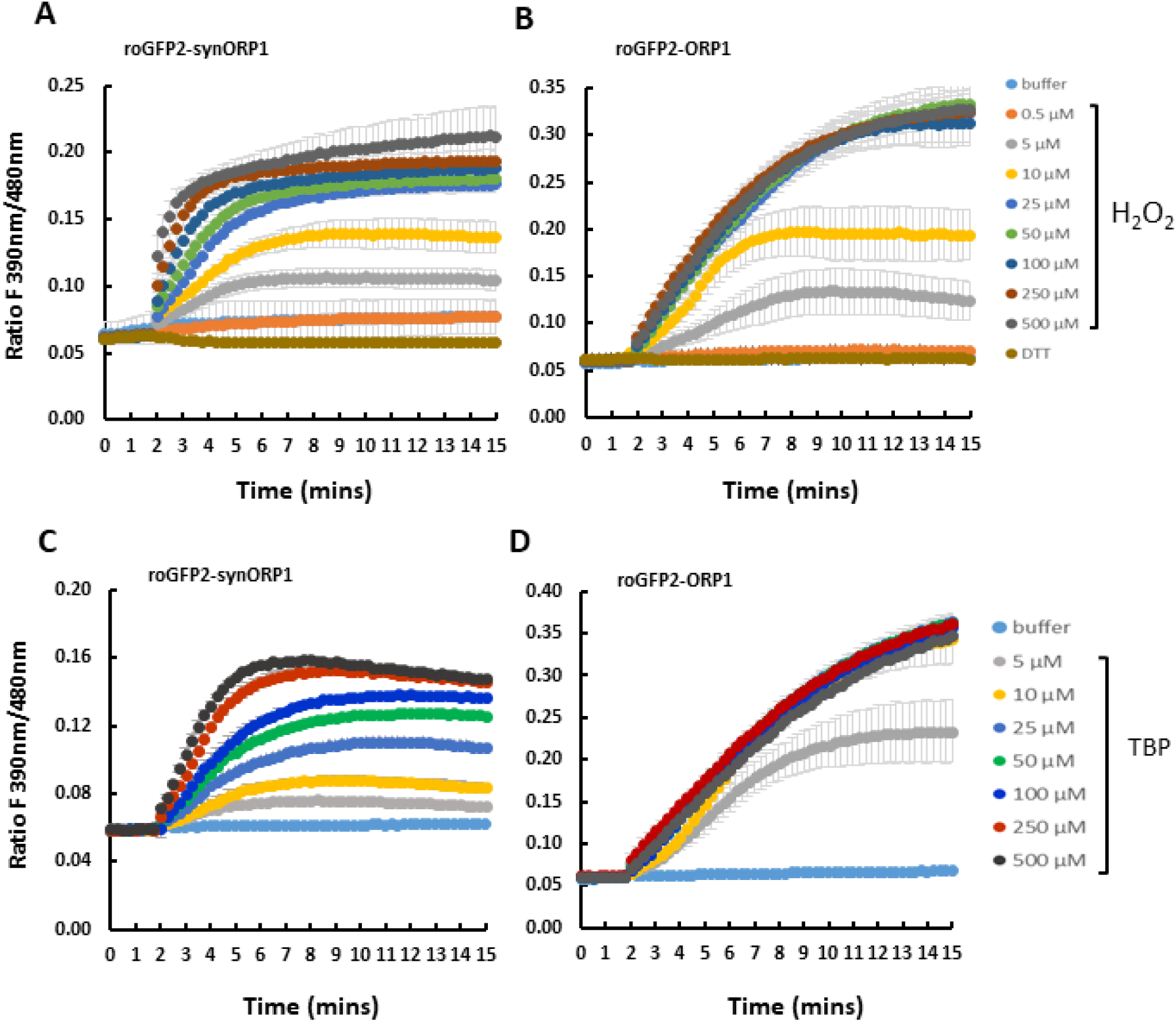
Oxidation of roGFP-(syn)ORP1 probes *in vitro* by H_2_O_2_ and TBP. **(A)** *O*xidation of 5 μM roGFP2-synORP1 or **(B)** roGFP2-ORP1 with 5 μM, 10 μM, 25 μM, 50 μM, 100 μM, 250 μM and 500 μM H_2_O_2_ or by the same concentrations of TBP **(C, D).** Each biosensor was also exposed to 1 mM DTT or buffer alone. Addition of peroxide or DTT was carried out 90 seconds after initiating measurements. Readings are means (± SD) combined from 4-6 technical replicates from two separate preparations.

**Supplemental Figure 4.**
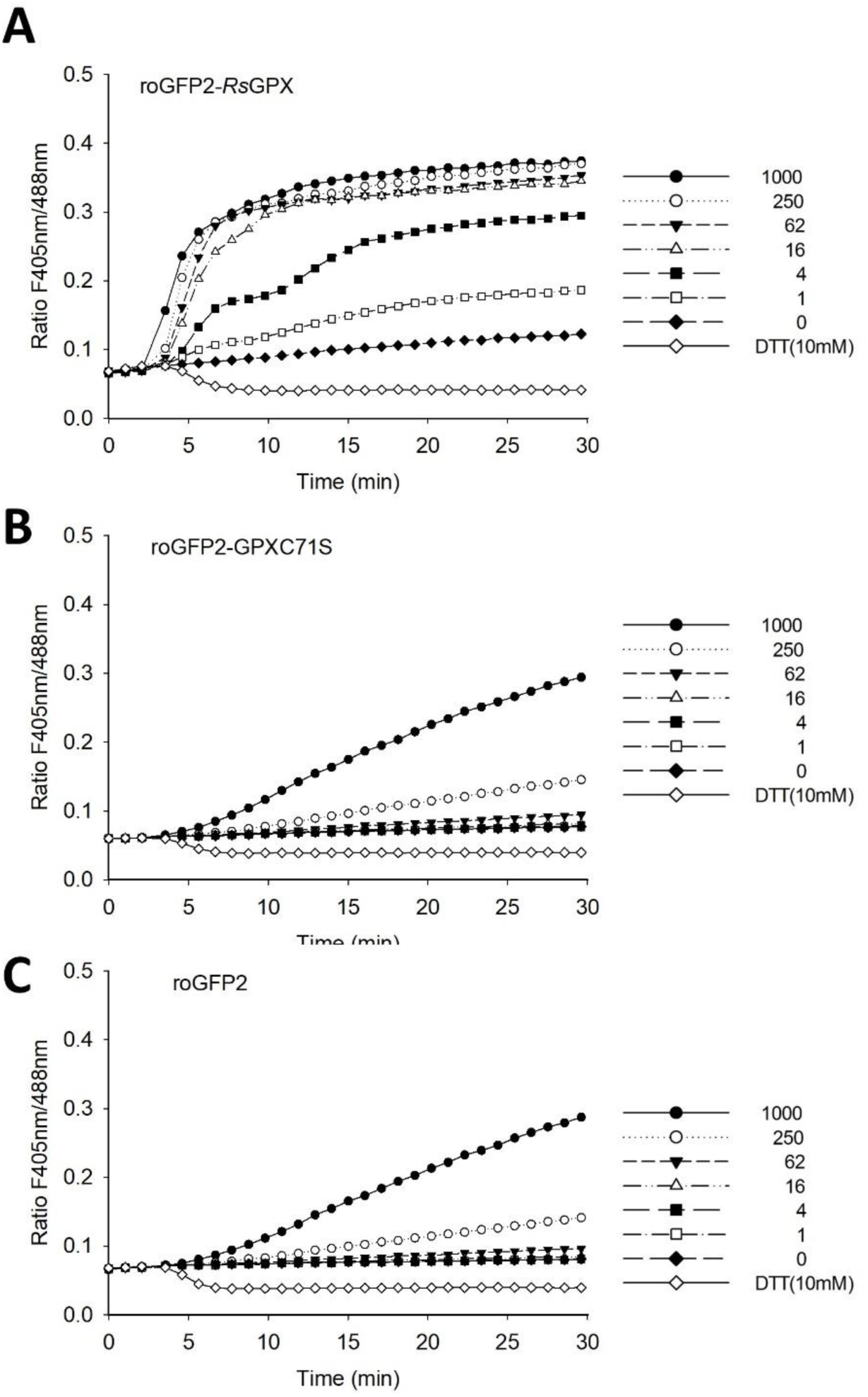
The mutation C71S on RsGPX abolishes the redox relay to roGFP2. **(A)** Response of recombinant reduced roGFP2-RsGPX **(B),** roGFP-GPXC71S **(C)** and roGFP2 to DTT and increasing concentration of H_2_O_2_ (1-1000µM). 5µM of each reduced biosensor was exposed to H_2_O_2_ 3 min after starting the measurement. The response of roGFP2 in the absence of any moiety was used as a negative control.

**Supplemental Figure 5.**
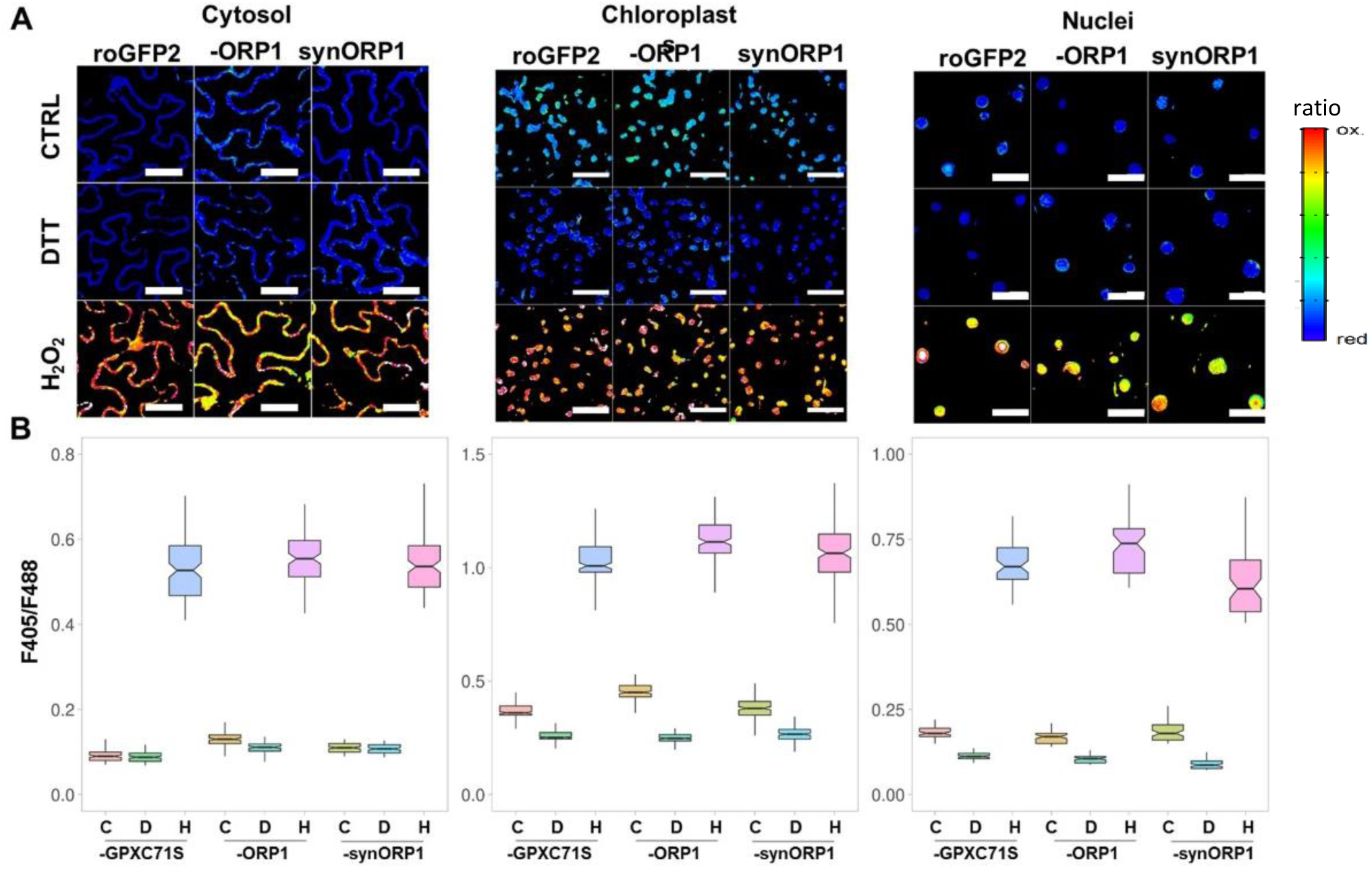
RoGFP2-ORP1, roGFP2-synORP1 and roGFP2-GPXC71S respond to extreme oxidation and reduction in different subcellular compartments of *Nicotiana benthamiana* abaxial epidermal cells. (A) Ratiometric images (F405/488 nm) of fluorescence emission at 530 nm reporting the oxidation state of the probes 5 min after infiltration of leaf discs with 100 mM H_2_O_2_ or 10 mM DTT treatment to achieve the maximum oxidation and reduction of biosensors, respectively. A water-only control treatment (CTRL) is shown for comparison. The constructs expressed biosensors in chloroplast stroma-, cytosol- and nuclei. Scale bar, 50 µm. **(B)** Quantified biosensor responses of roGFP2-GPXC71S, roGFP2-ORP1 and roGFP2-synORP1 in the three compartments. Box plots show responses to control (C, yellow), DTT (D, green) and H_2_O_2_ (H, blue/purple/pink) treatments. The Y axis shows the ratio (F405/488 nm) of fluorescence emission at 530 nm, reporting the oxidation state of the probes 5 min after infiltration of leaf discs. Within each box (interquartile range), the horizontal thick black line shows the median ratio for each condition, while indentations depict its 95% compatibility interval.

**Supplemental Data 1**. **Sequences of the roGFP2-(syn)ORP1 fusions inserted into pET28a(+).** The underlined letters mark the restriction enzymes sites for Nde I (5’ end) and Hind III (3’ end).

><u>NdeI</u> roGFP2 <u>HindIII</u>

<u>`CATATG`</u>GTGAGCAAGGGCGAGGAGCTGTTCACCGGGGTGGTGCCCATCCTGGTCGAGCTGGACGGCGACGTAAACGGCCACAAGTTCAGCGTGT CCGGCGAGGGCGAGGGCGATGCCACCTACGGCAAGCTGACCCTGAAGTTCATCTCCACCACCGGCAAGCTGCCCGTGCCCTGGCCCACCCTCGTGACCACCCTGACCTACGGCGTGCAGTGCTTCAGCCGCTACCCCGACCACATGAAGCAGCACGACTTCTTCAAGTCCGCCATGCCCGAAGGCTACGTCCAGGAGCGCACCATCTTCTTCAAGGACGACGGCAACTACAAGACCCGCGCCGAGGTGAAGTTCGAGGGCGACACCCTGGTGAACCGCATCGAGCTGAAGGGCATCGACTTCAAGGAGGACGGCAACATCCTGGGGCACAAGCTGGAGTACAACTACAACTGCCACAACGTCTATATCATGGCCGACAAGCAGAAGAACGGCATCAAGGTGAACTTCAAGATCCGCCACAACATCGAGGACGGCAGCGTGCAGCTCGCCGACCACTACCAGCAGAACACCCCCATCGGCGACGGCCCCGTGCTGCTGCCCGACAACCACTACCTGAGCACCTGCTCCGCCCTGAGCAAAGACCCCAACGAGAAGCGCGATCACA TGGTCCTGCTGGAGTTCGTGACCGCCGCCGGGATCACTCTCGGCATGGACGAGCTGTACAAGTAA<u>AAGCTT</u>

`>NdeI roGFP2-ORP1 HindIII`

<u>`CATATG`</u>GTTAGCAAAGGTGAAGAACTGTTCACCGGTGTTGTGCCGATCCTGGTTGAACTGGACGGTGACGTTAACGGTCACAAATTCTCTGTTT CCGGTGAAGGTGAAGGTGACGCTACCTACGGTAAACTGACCCTGAAATTCATCTCTACCACCGGTAAACTGCCGGTTCCGTGGCCGACCCTGGTGACCACCCTGACCTACGGTGTTCAGTGCTTCTCCCGTTACCCGGACCACATGAAACAGCACGATTTCTTCAAATCCGCGATGCCGGAAGGTTACGTGCAGGAACGTACCATCTTCTTCAAAGATGACGGTAACTACAAAACCCGTGCTGAAGTTAAATTCGAAGGTGACACCCTGGTTAACCGTATTGAACTGAAAGGCATCGACTTCAAAGAAGATGGTAACATCCTGGGTCACAAACTGGAATACAACTACAACTGTCACAACGTTTACATCATGGCGGATAAACAGAAAAACGGTATCAAAGTTAACTTCAAAATCCGTCACAACATCGAAGATGGTTCTGTTCAGCTGGCTGATCACTACCAGCAGAACACCCCGATCGGTGACGGTCCGGTTCTGCTGCCGGACAACCACTACCTGTCTACCTGCTCCGCGCTGAGCAAAGACCCGAACGAAAAACGTGACCACATGGTTCTGCTGGAATTCGTTACCGCGGCGGGTATCACCCTGGGCATGGATGAACTGTACAAAACCTCCGGCGGTTCTGGTGGCGGTGGCTCTGGTGGTGGTGGTTCTGGTGGTGGCGGCTCTGGTGGTGGTGGTTCTGGTGGCGGTGGTTCTGGCGGCGAATTCGACATCAGCGAATTCTACAAACTGGCTCCGGTTGACAAAAAAGGTCAGCCGTTCCCGTTCGACCAGCTGAAAGGTAAAGTGGTGCTGATCGTTAACGTTGCATCTAAATGCGGTTTCACCCCGCAGTACAAAGAACTGGAAGCGCTGTATAAACGTTACAAAGATGAAGGTTTCACTATCATCGGCTTCCCGTGTAACCAGTTCGGTCACCAGGAACCGGGCTCTGATGAAGAAATCgcCCAGTTCTGCCAACTGAACTATGGCGTGACTTTCCCGATCATGAAAAAGATCGACGTTAACGGTGGTAACGAAGATCCGGTTTATAAATTCCTGaaGAGCCAAAAATCCGGTATGTTGGGCTTGAGAGGTATCAAATGGAACTTCGAAAAATTCCTGGTTG ACAAAAAAGGTAAAGTTTACGAACGTTATAGCTCTCTGACCAAACCGTCCTCTCTGTCTGAAACCATCGAAGAACTGCTGAAAGAAGTTT<u>AAGC</u><u>`TT`</u>

`>NdeI roGFP2-synORP1 HindIII`

<u>`CATATG`</u>GTTAGCAAAGGTGAAGAACTGTTCACCGGTGTTGTGCCGATCCTGGTTGAACTGGACGGTGACGTTAACGGTCACAAATTCTCTGTTT CCGGTGAAGGTGAAGGTGACGCTACCTACGGTAAACTGACCCTGAAATTCATCTCTACCACCGGTAAACTGCCGGTTCCGTGGCCGACCCTGGTGACCACCCTGACCTACGGTGTTCAGTGCTTCTCCCGTTACCCGGACCACATGAAACAGCACGATTTCTTCAAATCCGCGATGCCGGAAGGTTACGTGCAGGAACGTACCATCTTCTTCAAAGATGACGGTAACTACAAAACCCGTGCTGAAGTTAAATTCGAAGGTGACACCCTGGTTAACCGTATTGAACTGAAAGGCATCGACTTCAAAGAAGATGGTAACATCCTGGGTCACAAACTGGAATACAACTACAACTGTCACAACGTTTACATCATGGCGGATAAACAGAAAAACGGTATCAAAGTTAACTTCAAAATCCGTCACAACATCGAAGATGGTTCTGTTCAGCTGGCTGATCACTACCAGCAGAACACCCCGATCGGTGACGGTCCGGTTCTGCTGCCGGACAACCACTACCTGTCTACCTGCTCCGCGCTGAGCAAAGACCCGAACGAAAAACGTGACCACATGGTTCTGCTGGAATTCGTTACCGCGGCGGGTATCACCCTGGGCATGGATGAACTGTACAAAACCTCCGGCGGTTCTGGTGGCGGTGGCTCTGGTGGTGGTGGTTCTGGTGGTGGCGGCTCTGGTGGTGGTGGTTCTGGTGGCGGTGGTTCTGGCGGCGAATTCGACATCAGCGAATTCTACAAACTGGCTCCGGTTGACAAAAAAGGTCAGCCGTTCCCGTTCGACCAGCTGAAAGGTAAAGTGGTGCTGATCGTTAACGTTGCATCTAAATGCGGTTTCACCCCGCAGTACAAAGAACTGGAAGCGCTGTATAAACGTTACAAAGATGAAGGTTTCACTATCATCGGCTTCCCGTGTAACCAGTTCGGTCACCAGGAACCGGGCTCTGATGAAGAAATCCTGAACTCTCTGAAATACGTTCGTCCGGGTGGTGGTTTCGAACCGAACTTCCCGATCATGAAAAAGATCGACGTTAACGGTGGTAACGAAGATCCGGTTTATAAATTCCTGCGTGAAGCGCTGCCGGCTCCGTCTGATGACGCGACCGCGCTGATGACCGATCCGAAACTGATCACCTGGAGCCCGGTTTGCCGTAACGATGTTGCGTGGAACTTCGAAAAATTCCTGGTTGACAAAAAAGGTAAAGTTTACGAACG TTATAGCTCTCTGACCAAACCGTCCTCTCTGTCTGAAACCATCGAAGAACTGCTGAAAGAAGTTGAATAA<u>AAGCTT</u>

`>NdeI roGFP2-GPXC71S HindIII`

<u>`CATATG`</u>GTTTCTAAAGGTGAAGAACTGTTCACCGGTGTTGTTCCGATCCTGGTTGAACTGGATGGTGACGTTAACGGTCACAAATTCTCTGTTT CTGGTGAAGGTGAAGGTGATGCTACCTACGGTAAACTGACCCTGAAATTCATCTCCACCACTGGTAAACTGCCGGTGCCGTGGCCGACCCTGGTTACCACCCTGACCTACGGCGTTCAGTGCTTCTCTCGTTACCCGGACCACATGAAACAGCACGACTTCTTCAAATCCGCTATGCCGGAAGGCTACGTTCAGGAACGCACCATTTTCTTCAAAGATGACGGTAACTACAAAACCCGCGCTGAAGTTAAATTCGAAGGTGATACTCTGGTTAACCGTATCGAACTGAAAGGTATCGATTTCAAAGAAGACGGTAACATCCTGGGTCACAAACTGGAATACAACTACAACTGCCACAACGTGTACATCATGGCGGACAAACAGAAAAACGGTATTAAAGTTAACTTCAAAATCCGTCACAACATCGAAGATGGCAGCGTGCAGCTGGCCGACCACTACCAGCAGAACACCCCGATCGGTGACGGTCCGGTTCTGCTGCCGGACAACCACTACCTGTCCACCTGCTCCGCGCTGTCTAAAGACCCGAACGAAAAACGTGATCACATGGTTCTGCTGGAATTCGTTACCGCAGCGGGCATCACCCTGGGTATGGACGAACTGTACAAAACTTCTGGTGGTTCTGGCGGTGGCGGCTCTGGCGGTGGTGGTAGCGGCGGCGGTGGTTCTGGTGGCGGTGGTTCCGGTGGCGGCGGTAGCGGTGGCGAATTCGACCAGTCTTCTTACTCTTCCATCTACCACATCTCTGTTAAAGACATCGATGGCAACGACGTGTCTCTGTCTAAATTCACCGGTAAAGTGCTGCTGATCGTAAACGTTGCGTCTAAATCTGGTCTGACCCAGGGTAACTACAAAGAACTGAACATCCTGTACGCGAAATACAAAACCAAAGGTCTGGAAATCCTGGCGTTCCCGTGTAACCAGTTCGGTTCTCAGGAACCGGGTTCTAACAAAGAAATTAAAGATAACATCTGTACCACCTTTAAAGGTGAATTCCCGATCTTTGATAAAATTGAGGTTAACGGTGAAAACGCGTCCCCACTGTATAAATTTCTGAAAGAGCAGAAAGGCGGTCTGTTCGGTGATAGCATTAAATGGAACTTTGCTAAATTCTTAGTTGATAAACAAGGTAACGTAGTTGATCGTTTCGCGCCGACCACTAGCCCGTTAGAGATTGAGAAAGACATTGAAAAACTGCTGGCATC TACCTAA<u>AAGCTT</u>

